# TNF signaling mediates cellular immune function and promotes malaria parasite killing in the mosquito *Anopheles gambiae*

**DOI:** 10.1101/2024.05.02.592209

**Authors:** George-Rafael Samantsidis, Hyeogsun Kwon, Megan Wendland, Catherine Fonder, Ryan C. Smith

## Abstract

Tumor Necrosis Factor-α (TNF-α) is a proinflammatory cytokine and a master regulator of immune cell function in vertebrates. While previous studies have implicated TNF signaling in invertebrate immunity, the roles of TNF in mosquito innate immunity and vector competence have yet to be explored. Herein, we confirm the identification of a conserved TNF-α pathway in *Anopheles gambiae* consisting of the TNF-α ligand, Eiger, and its cognate receptors Wengen and Grindelwald. Through gene expression analysis, RNAi, and *in vivo* injection of recombinant TNF-α, we provide direct evidence for the requirement of TNF signaling in regulating mosquito immune cell function by promoting granulocyte midgut attachment, increased granulocyte abundance, and oenocytoid rupture. Moreover, our data demonstrate that TNF signaling is an integral component of anti-*Plasmodium* immunity that limits malaria parasite survival. Together, our data support the existence of a highly conserved TNF signaling pathway in mosquitoes that mediates cellular immunity and influences *Plasmodium* infection outcomes, offering potential new approaches to interfere with malaria transmission by targeting the mosquito host.

## Introduction

*Anopheles* mosquitoes serve as the primary vectors of *Plasmodium* parasites, which cause malaria and impose substantial burdens on public health across the globe [1]. While there has been a significant reduction in malaria cases over the last twenty years due to improved vector control strategies, the continued effectiveness of these strategies has been jeopardized by increased insecticide resistance [2–4], which highlights the need to develop alternative approaches for malaria control. With recent advancements in gene-drive systems offering significant promise for population modification [5–7], the potential that these genetic techniques can be used to manipulate the vector competence of mosquito populations to impair malaria transmission has become a reality. However, to fully leverage these genetic approaches, we require a better understanding of the molecular mechanisms that define *Plasmodium* infection in the mosquito host.

In response to *Plasmodium* infection, mosquitoes mount a series of sequential immune signals initiated by the midgut in response to ookinete invasion [8–12], which are further processed by the mosquito immune cells (hemocytes) [13–15] to promote malaria parasite killing [16–19]. As a result, hemocytes serve as integral immune mediators that directly contribute to ookinete recognition [16,17] or that promote humoral responses to limit oocyst survival [15,17,19]. Mosquito hemocytes have traditionally been classified into three main cell types based on morphological and biochemical properties: granulocytes, oenocytoids, and prohemocytes [20], with more recent single-cell studies expanding on the complexity of these cell populations [21,22]. Macrophage-like granulocytes are phagocytic and behave as immune sentinels either in circulation in the hemolymph or as sessile cells attached to the midgut or other mosquito tissues [20,23]. Previous studies have demonstrated that granulocytes respond to stimuli resulting from ookinete midgut invasion to mediate both early- and late-phase immune responses against *Plasmodium* [15–18]. Oenocytoids have primarily been associated with the expression of prophenoloxidases (PPOs) [20], which are key enzymes in the melanization pathway and have been previously implicated in oocyst survival [19]. Lastly, prohemocytes are presumed precursors [20,24] that give rise to granulocyte and oenocytoid populations under infective conditions [13–15].

Mosquito hemocyte populations are highly heterogenic [21,22], with their composition tightly regulated by a variety of signaling pathways in response to different physiological conditions. This includes previous studies that have demonstrated the ability of hemocytes to proliferate in response to blood-feeding [24,25], a process regulated by the release of insulin-like peptides and subsequent activation of the PI3K/AKT and MAPK/ERK signaling pathways [26–29]. In addition, the Signal Transducer and Activator of Transcription (STAT) [14,15], the LPS-induced TNF-alpha factor (LITAF)-like transcription factor 3 (LL3) [15,30], c-Jun N-terminal kinase (JNK) [14], and Toll [14,30] pathways have been associated with hemocyte differentiation and parasite attrition. Furthermore, eicosanoid signaling pathways have been implicated in hemocyte function, differentiation, and *Plasmodium* killing [18,19,31,32], and are central to the establishment of innate immune memory that confers increased resistance to infection [18,31]. Yet, despite these advances, our understanding of the immune signals that modulate mosquito hemocytes remains limited.

Tumor Necrosis Factor-α (TNF-α) is one of the most important regulators of immune function in vertebrates, acting as a proinflammatory cytokine and critical mediator of immune cell regulation [33,34]. Across vertebrate systems, components of TNF signaling have remained conserved, consisting of a TNF-α ligand and two receptors: the TNFR1 and TNFR2 [34]. Previous phylogenetic analyses have revealed the presence of orthologous TNF pathways in invertebrates [35,36], however few studies have examined TNF signaling beyond *Drosophila*. With the *Drosophila* TNF pathway comprising an analogous TNF ligand, *Eiger*, and the receptors, *Wengen* (*Wgn*) and *Grindelwald* (*Grnd*) [37], *Drosophila* TNF signaling regulates several physiological processes, including tissue growth regulation, cellular proliferation, development, and host defense [38], with significant contributions of hemocytes in mediating these functions [39,40]. Under both homeostatic or infected conditions, Eiger modulates *Drosophila* hemocyte function to promote survival through actions as a chemoattractant [41], inducer of cell death [42], or regulator of phagocytosis [43–45]. While recent studies have expanded our knowledge on how TNF signaling influences the function of immune cells in other insect species [46], the role of TNF signaling in the mosquito innate immune system remains unclear.

In this study, we establish integral roles of mosquito TNF signaling in mediating anti-*Plasmodium* immune responses that limit malaria parasite survival in the mosquito host. While gene expression analysis indicates that *Eiger* is induced in response to blood-feeding regardless of infection status in midgut and hemocyte tissues, downstream experiments clearly demonstrate the roles of mosquito TNF signaling in cellular immune function and immune responses that promote malaria parasite killing. Together, our data provide novel mechanistic insight into the function of TNF signaling in mosquito immune cell regulation and anti-*Plasmodium* immunity.

## Results

### Expression analyses of mosquito TNF signaling pathway components

To better understand TNF signaling in *Anopheles gambiae* we examined the expression of the TNF ligand, *Eiger*, and two TNF receptors, *Wgn* and *Grnd* (**Fig 1A**), across tissues (midgut, hemocytes, and carcass) and physiological conditions (naïve, blood-fed, and *Plasmodium berghei* infection). Under naïve conditions, *Eiger* displayed comparable expression levels between hemocytes and the carcass, with reduced levels of expression in the midgut (**Fig 1B**). A similar expression pattern was observed for *Wgn*, although higher levels of *Wgn* expression were found in carcass tissues (**Fig 1C**). In contrast, *Grnd* was enriched in both midgut and carcass, with the lowest expression in hemocytes (**Fig 1D**). Of note, *Eiger* expression was generally increased across tissues in response to blood-feeding and infection, and was significantly induced in hemocytes regardless of infection status (**Fig 1B**). In contrast, the different feeding conditions had little effect on *Wgn* and *Grnd* expression, although *Wgn* displayed significantly reduced levels of expression in the carcass following *P. berghei* infection (**Fig 1C**), suggesting the potential down-regulation of TNF signaling in the carcass following infection. While both TNF and TNF receptors (TNFRs) are influenced by posttranscriptional modifications in *Drosophila* [47–49] and other vertebrate systems, the observed patterns of *Eiger* expression support potential roles of TNF signaling in mosquito immunity and cellular immune function.

**Figure 1.**
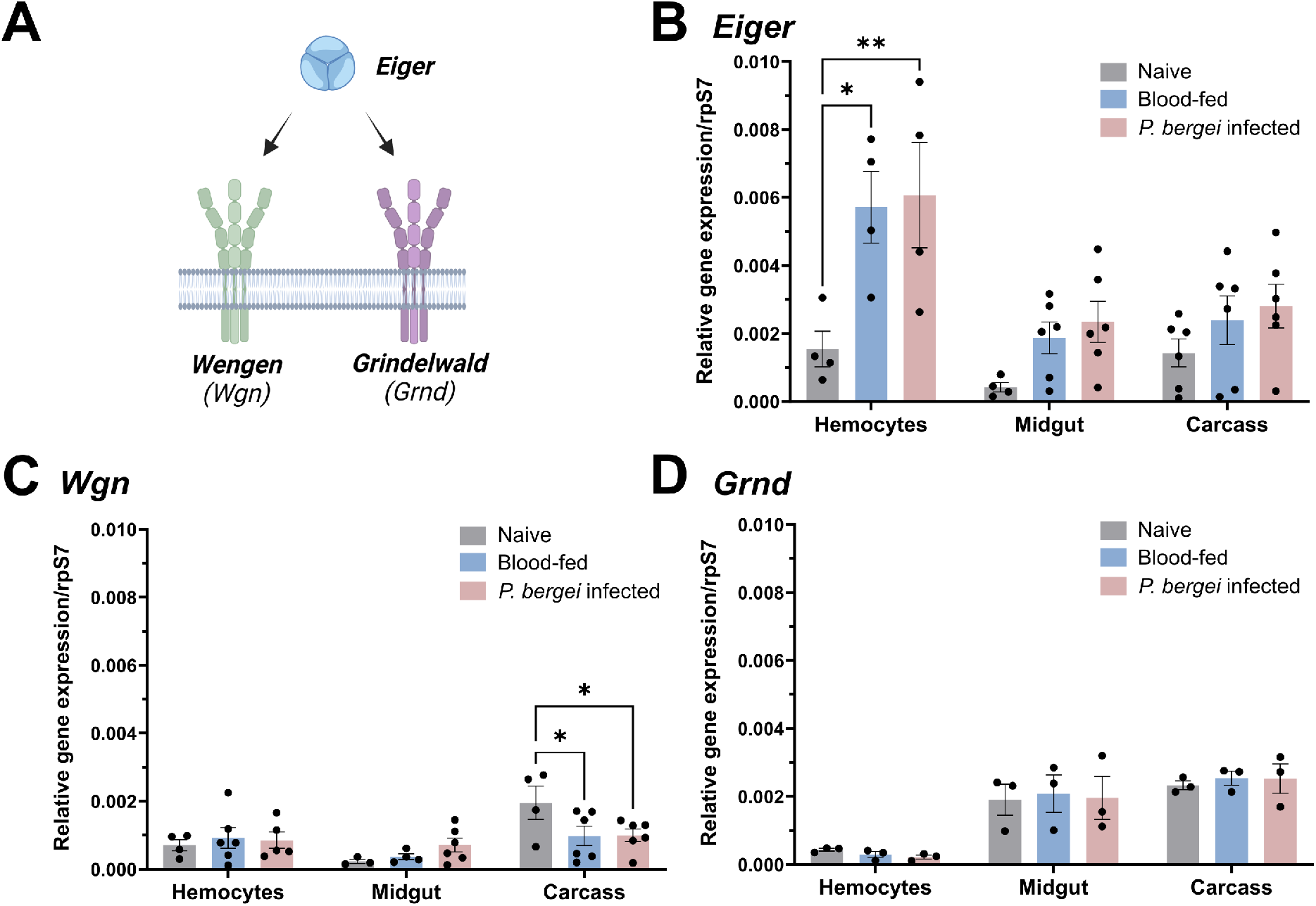
Expression patterns of *Eiger*, *Wgn*, and *Grnd* in mosquitoes. (**A**) Schematic representation of the mosquito TNF signaling pathway. The expression of *Eiger* (**B**)*, Wgn* (**C**), and *Grnd* (**D**) was examined by qPCR in the mosquito midgut, hemolymph, and fat body under naive, blood-fed (24h post-feeding) or *P. berghei-*infected (24h post-infection) conditions. Expression data are displayed relative to rpS7 expression with bars representing the mean ± SE of three to six independent biological replicates (black dots). Data were analyzed using a two-way ANOVA with a Tukey’s multiple comparisons test to determine significance. Asterisks denote significance (*P < 0.05, **P < 0.01).

### Mosquito TNF signaling limits *P. berghei* development

To examine the influence of TNF signaling on malaria parasite infection, we first injected adult female mosquitoes with 50ng of recombinant human TNF-α (rTNF-α) one day before challenging with *P. berghei*. When malaria parasite numbers were evaluated 8 days post-infection, mosquitoes primed with rTNF-α significantly reduced *P. berghei* oocyst numbers and the prevalence of infection (**Fig 2A**). Conversely, when *Eiger* is silenced via RNAi (**Fig S1**), oocyst numbers and infection prevalence were significantly increased (**Fig 2B**). Similarly, when the TNFRs *Wgn* and *Grnd* are silenced following the injection of dsRNA (**Fig S1**), both *Wgn*- and *Grnd*-silenced backgrounds resulted in significant increases in *P. berghei* oocyst numbers (**Fig 2C and 2D**). Together, these findings demonstrate the importance of mosquito TNF signaling in the innate immune response against *Plasmodium* and confirm the integral roles of Eiger, Wgn, and Grnd in this process.

**Figure 2.**
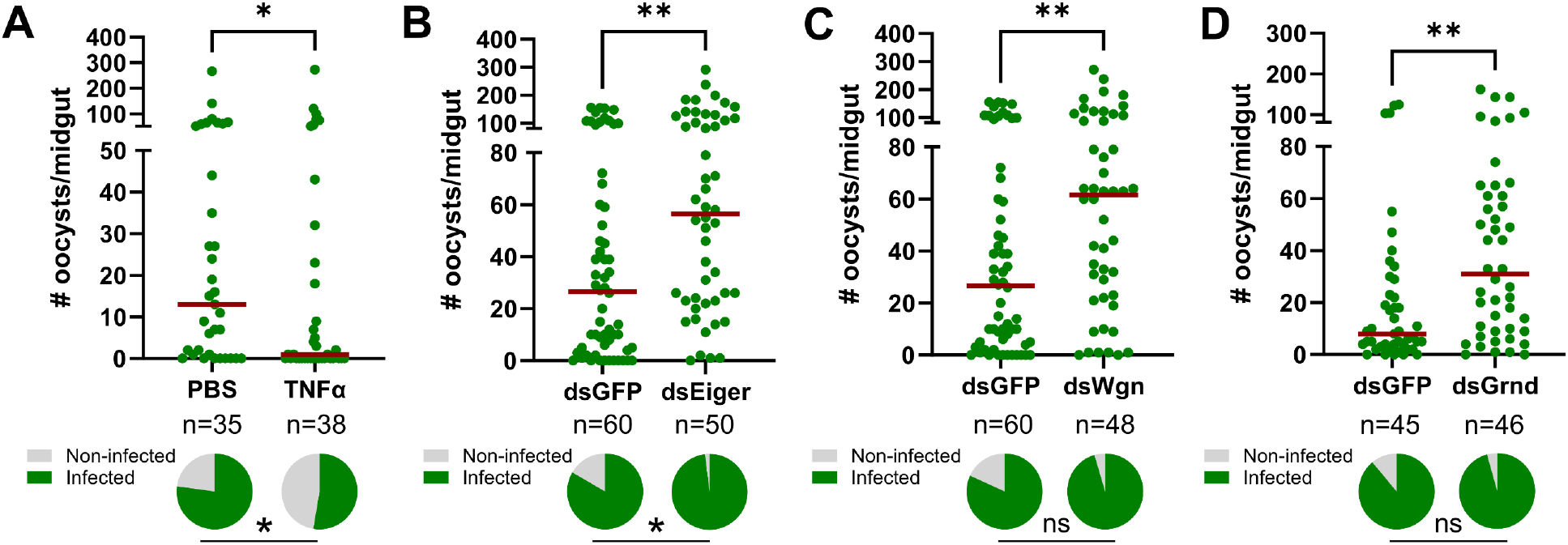
TNF signaling in *An*. *gambiae* limits *Plasmodium* survival. (**A**) Adult female mosquitoes were injected with 1XPBS (control) or 50ng of rTNF-α prior to infection with *P. berghei.* Oocyst numbers and infection prevalence were evaluated at 8 days post infection. Additional RNAi experiments were performed to evaluate the contributions of the TNF signaling components *Eiger* (**B**), *Wgn* (**C**), and *Grnd* (**D**) in the context of *P. berghei* infection. Oocyst numbers and infection prevalence were similarly evaluated at 8 days post infection. Mosquitoes injected with dsGFP served as control in all experiments. For each graph, dots correspond to the number of oocysts identified in individual midguts, with the median represented by a red horizontal line. Infection prevalence (% infected/total) is depicted as pie charts pies below each figure. Data were combined from three or more independent experiments. Statistical significance was determined using a Mann-Whitney test to assess oocyst numbers, while a Fisher’s Exact test was performed to measure differences in infection prevalence. Asterisks denote statistical significance (* *P* < 0.05, \*\**P* < 0.01). n= numbers of individual mosquitoes examined.

### Wgn and Grnd comprise a singular pathway to promote anti-*Plasmodium* immunity

In vertebrate systems, TNF signaling can initiate distinct cellular responses based on the interactions of TNF-α with its cognate TNFR1 and TNFR2 receptors [33,34]. With both Wgn and Grnd serving as antagonists to *Plasmodium* development (**Fig 2**), we wanted to examine whether TNF signaling via Wgn or Grnd produced dependent or independent responses that contribute to parasite killing. Similar to our results in **Fig 2C** and **2D**, the silencing of *Wgn* or *Grnd* resulted in increased *P. berghei* oocyst numbers (**Fig 3**). However, when both *Wgn* and *Grnd* were silenced, the infection intensity did not further increase (**Fig 3**). This suggests that *Wgn* and *Grnd* work together in a singular pathway to promote malaria parasite killing, where the loss of either component abrogates TNF signaling.

**Figure 3.**
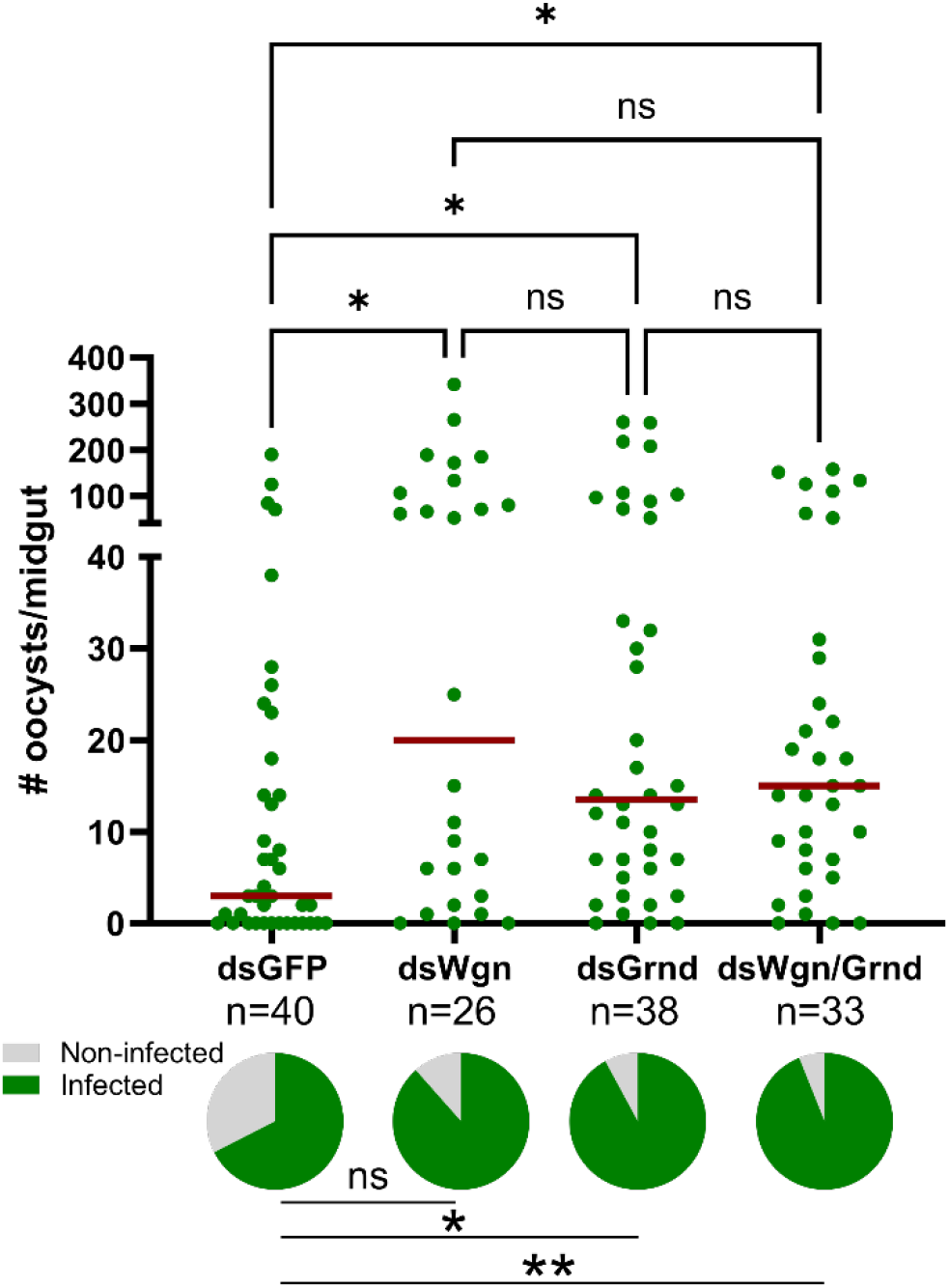
TNF signaling requires the concerted function of Wgn and Grnd to promote parasite killing. RNAi experiments were performed to evaluate the contributions of the TNF signaling components Eiger, Wgn, and Grnd in the context of *P. berghei* infection. Oocyst numbers and infection prevalence were evaluated at 8 days post-infection in *GFP*-, *Wgn*-, *Grnd*-, and *Wgn/Grnd*-silenced backgrounds. For each graph, dots correspond to the number of oocysts identified in individual midguts, with the median represented by a red horizontal line. Infection prevalence (% infected/total) is depicted as pie charts pies below each figure. Data were combined from three or more independent experiments. Statistical significance was determined using Kruskal-Wallis with a Dunn’s multiple comparison test to assess oocyst numbers, while a Fisher’s Exact test was performed to measure differences in infection prevalence. Asterisks denote statistical significance (* *P* < 0.05, \*\**P* < 0.01). ns, not significant; n= numbers of individual mosquitoes examined.

### Mosquito TNF signaling modulates granulocyte and oenocytoid populations

Previous studies have demonstrated that mosquito hemocyte populations undergo significant changes in response to different physiological conditions [13–19,24,25,29] and have established their significant roles in malaria parasite killing [14–19]. Based on the upregulation of *Eiger* in perfused hemocyte samples (**Fig 1**) and the importance of TNF signaling associated with immune cell regulation in other systems [33,34,42,46], we wanted to explore the effects of TNF signaling on mosquito hemocyte populations.

With functional activation of the TNF pathway requiring TNF-α to initiate signaling via its cognate receptors, the expression patterns of *Wgn* and *Grnd* were first examined using previous single-cell transcriptomic data of mosquito hemocytes [22]. While *Wgn* is enriched in both granulocyte and oenocytoid populations, *Grnd* is only expressed in oenocytoids (**Fig 4A** and **4C**, **Fig S2**). Additional experiments using clodronate liposomes to deplete phagocytic granulocyte populations [17,22,50,51] further support the enrichment of *Wgn* in granulocytes, where granulocyte depletion resulted in a significant reduction in *Wgn* expression (**Fig 4B**). In contrast, *Grnd* expression increased following granulocyte depletion, which is likely the result of the enrichment of non-phagocytic oenocytoid populations (**Fig 4C**).

**Figure 4.**
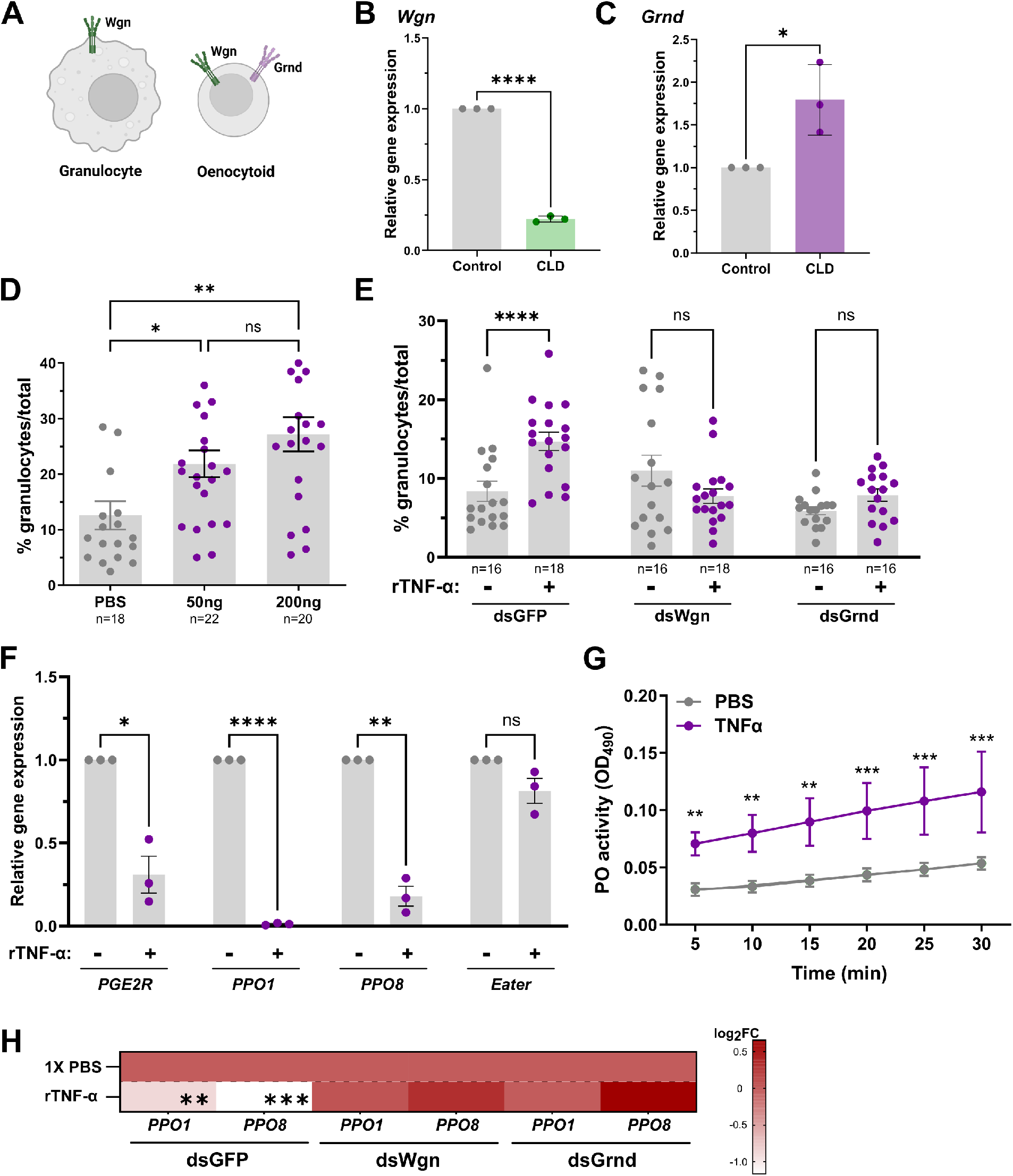
Expression of *Wgn* and *Grnd* in mosquito hemocytes. (**A**) Previous single cell transcriptomic data [22] support the expression of *Wgn* in both granulocytes and oenocytoids, whereas *Grnd* is enriched specifically in oenocytoids. The expression levels of *Wgn* (**B**) and *Grnd* (**C**) were examined in mosquitoes treated with either clodronate liposomes (CLD) to deplete mosquito granulocytes or control liposomes to functionally demonstrate the specificity of *Grnd* to oenocytoids using qPCR. Data from three independent experiments were analyzed by an unpaired Students’ t-test. (**D**) Injection with rTNF-α (50ng or 200ng) increases the granulocyte proportions at 24h post-injection compared to 1XPBS controls. Similar experiments performed in *GFP*-, *Wgn*-, and *Grnd*-silenced backgrounds demonstrate the importance of both *Wgn* and *Grnd* for the increase in granulocytes following rTNF-α (50ng) injection (**E**). For both **D** and **E**, the percentage of granulocytes of the total hemocytes are represented as mean ± SE of three independent biological replicates with statistical significance determined by Mann-Whitney to compare the effects of rTNF-α versus the 1XPBS control mosquitoes. (**F**) Injection of mosquitoes with TNF-α reduces the expression of oenocytoid specific genes (*PPO1, PPO8,* and *PGE2R)*, suggesting that TNF promotes oenocytoid lysis. The granulocyte marker *Eater* was used as a negative control. Data from three independent experiments were analyzed by an unpaired Students’ t-test. (**G**) Additional experiments were performed to determine the effects of rTNF-α on the phenoloxidase (PO)-activity of mosquito hemolymph (n=20). Six measurements (OD490) were taken for DOPA conversion assays at 5-min intervals. Bars represent mean ± SE of three independent biological experiments with statistical significance determined with a two-way repeated-measures ANOVA followed by Sidak’s multiple comparison test. (**H**) Silencing the expression of *Wgn* and *Grnd* impaired the rTNF-α induced phenotypes on *PPO1* and *PPO8* expression when tested in whole adult mosquitoes. Results are presented as a heatmap displaying the log_2_ fold change (FC) and indicate differences gene expression as measured by qPCR following treatment with rTNF-α or 1X-PBS (control). Data represent the mean fold change expression of three independent biological replicates, with significance determined using an unpaired Students’ t-test. Asterisks indicate significance (* *P* < 0.05, \*\**P* < 0.01, **** *P* < 0.0001). ns, not significant; n= numbers of individual mosquitoes examined.

To determine the effects of mosquito TNF signaling on hemocyte subpopulations, we first injected mosquitoes with rTNF-α and examined the effects on granulocyte numbers. The injection of mosquitoes with either 50ng or 200ng of rTNF-α caused a substantial increase in the proportion of granulocytes at comparable levels (**Fig 4D**). Based on patterns of expression in hemocyte subtypes (**Fig 4A**, **Fig S2**), we hypothesized that rTNF-α would likely influence granulocyte populations via Wgn signaling. While rTNF-α similarly increased the percentage of granulocytes in the control *GFP*-silenced background, *Wgn*-silencing negated the effects of rTNF-α on granulocyte proportions as expected (**Fig 4E**). Interestingly, the same effect was also observed following rTNF-α treatment in *Grnd*-silenced mosquitoes (**Fig 4E**), suggesting that both *Wgn* and *Grnd* are required for the rTNF-α-mediated increase in the percentage of granulocytes. Given the absence of *Grnd* in granulocytes (**Fig 4A**, **Fig S2**), there is support for an indirect model in which the effects of rTNF-α on granulocytes are mediated by oenocytoid function.

With previous studies in *Drosophila* demonstrating that TNF signaling promotes melanization through crystal cell lysis [42,44], the equivalent of mosquito oenocytoid immune cell populations, we wanted to explore if TNF signaling similarly promotes oenocytoid lysis. To address this question, we injected mosquitoes with rTNF-α and analyzed the expression levels of *PPO1*, *PPO8,* and *PGE2R*, genes enriched in oenocytoids [19,22], which have previously been used to illustrate oenocytoid cell rupture [19]. At 24hrs post-injection*, PPO1, PPO8,* and *PGE2R* each displayed significantly reduced gene expression when compared to controls (**Fig 4F**), indicative of oenocytoid lysis. In contrast, the expression of the granulocyte marker, *Eater,* remained unchanged (**Fig 4F**). With the rupture of oenocytoids releasing pro-phenoloxidases and other cellular contents into the hemolymph, we performed dopa conversion assays to measure hemolymph phenoloxidase (PO) activity following the injection of rTNF-α as an additional measurement of oenocytoid rupture [19]. As expected, TNF-α treatment significantly increased hemolymph PO activity when compared to control mosquitoes (**Fig 4G**), providing further evidence that TNF signaling promotes oenocytoid lysis/rupture. Additional experiments with rTNF-α in *Grnd-* or *Wgn*-silenced backgrounds negated the rTNF-α-mediated down-regulation of *PPO1* and *PPO8* expression (**Fig 4H**), suggesting that both *Wgn* and *Grnd* are required to promote oenocytoid lysis via mosquito TNF signaling. When paired with the requirement of both receptors to promote parasite killing (**Fig 3**), these data provide further support that *Wgn* and *Grnd* act together to initiate the TNF-mediated signals that promote oenocytoid lysis.

### TNF-mediated *Plasmodium* killing is independent of granulocyte function

Previous studies have shown that both granulocytes and oenocytoids have central roles in anti-*Plasmodium* immunity [17,19,52], which given the effects of TNF signaling on granulocyte numbers and oenocytoid lysis (**Fig 4**), may corroborate our observations regarding the influence of mosquito TNF signaling on *Plasmodium* infection (**Figs 2 and 3**). Since granulocytes play pivotal roles in ookinete recognition [16,17], we examined the effects of TNF signaling on early oocyst numbers. Similar to the results presented in **Fig 2A**, rTNF-α injection significantly reduced *P. berghei* early oocyst numbers when observed at 2 days post-infection (**Fig 5A**), suggesting that TNF signaling may enhance ookinete recognition. This is further supported by the increased attachment of mosquito hemocytes to the mosquito midgut following rTNF-α injection (**Fig 5B**), which suggests that TNF signaling contributes to early-phase anti-*Plasmodium* killing responses. To further confirm the role of granulocytes in TNF-mediated parasite killing, we again employed the use of clodronate liposomes to deplete mosquito granulocyte populations [17,22,50,51]. To approach this question, we first depleted phagocytic granulocytes using clodronate liposomes, then treated mosquitoes with rTNF-α prior to challenge with *P. berghei* (**Fig 5C**). Before infection experiments, hemolymph was perfused and the granulocyte proportions were examined under each experimental condition to confirm granulocyte depletion. Similar to **Fig 4D**, the injection of rTNF-α in the control liposome background resulted in increased granulocyte numbers (**Fig 5D**). Additional experiments following clodronate liposome treatment confirm the successful depletion of granulocytes independent of rTNF-α treatment (**Fig 5B**), thereby providing a methodology to examine the TNF-mediated contributions of granulocytes to anti-*Plasmodium* immunity. While TNF-α injection reduced the parasite load in mosquitoes treated with control liposomes (**Fig 5E**) similar to previous experiments (**Fig 2A** and **5A**), *P. berghei* oocyst numbers were significantly increased in PBS treated mosquitoes in the granulocyte-depleted background (**Fig 5E**) as previously reported [17]. Of note, when rTNF-α injection was performed in the clodronate liposome background, parasite numbers were significantly reduced (**Fig 5E**), suggesting that granulocyte depletion does not fully impair the TNF-mediated mechanisms that promote parasite killing. As a result, other TNF-mediated effects on oenocytoid function that influence hemolymph PO activity (**Fig 4**) and oocyst killing responses [17,19] may also contribute to malaria parasite killing (**Fig 6**). However, we cannot rule out the potential effects of TNF signaling on humoral immune responses produced by the fat body.

**Figure 5.**
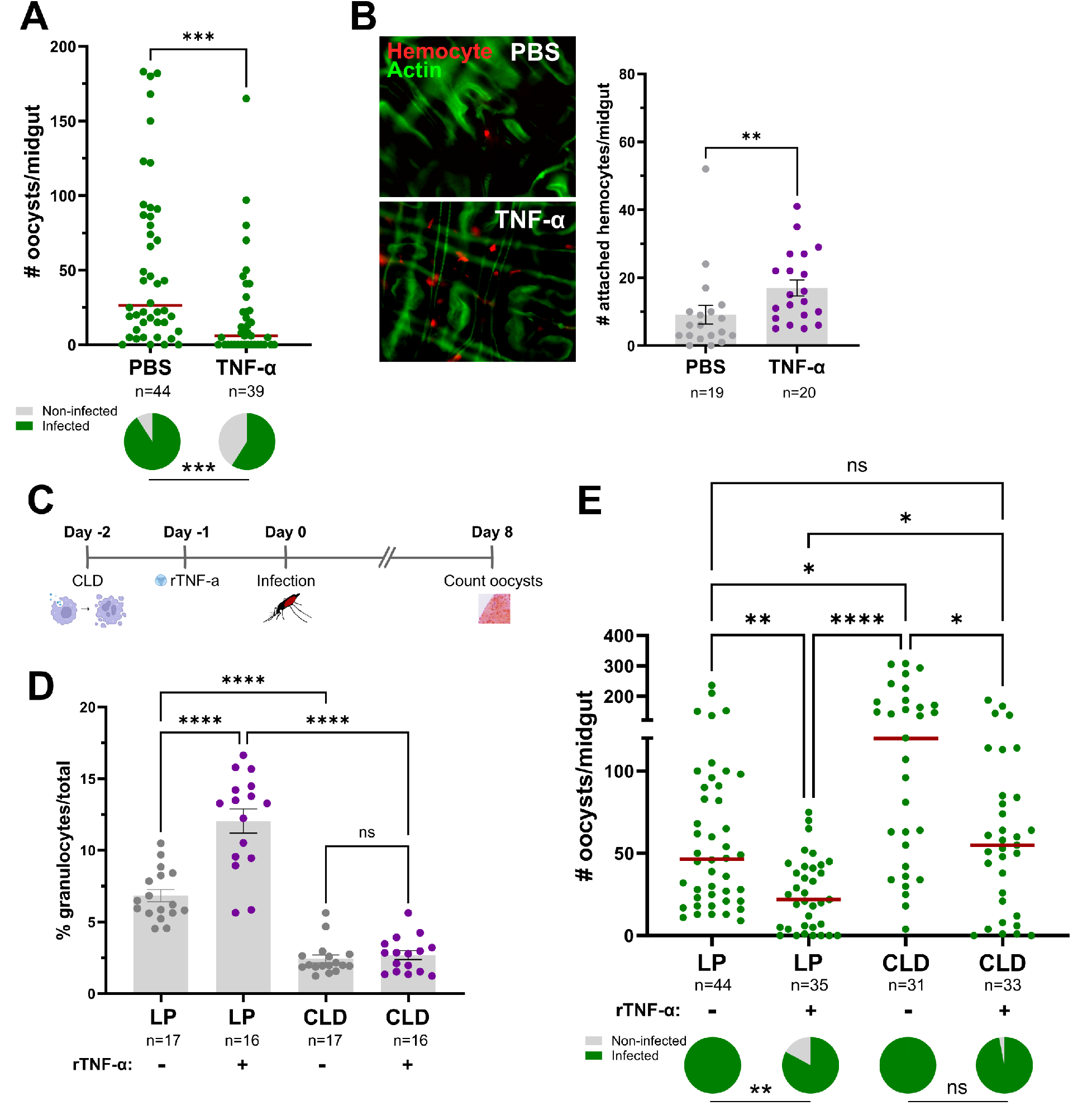
TNF-mediated parasite killing targets ookinete invasion and is mediated in part by granulocyte function. (**A**) Adult female mosquitoes were injected with 1XPBS (control) or 50ng of rTNF-α prior to infection with *P. berghei*. Oocyst numbers and infection prevalence were evaluated at 2 days post infection to measure early oocyst numbers and the success of ookinete invasion. (**B**) Immunofluorescence images of DiI-stained hemocytes (red) attached to the mosquito midgut (counterstained with phalloidin, green) approximately 24 hrs post-treatment with 1xPBS (control) or 50ng of rTNF-α. The number of attached hemocytes was quantified for each respective treatment with the dots corresponding to the number of hemocytes attached to each individual midgut examined. To address the role of granulocytes in TNF-mediated parasite killing, mosquitoes were first injected with either clodronate liposomes (CLD) to deplete granulocytes or control liposomes (LP), then 24 hours later surviving mosquitoes were treated with 50ng rTNF-α or 1X-PBS (**C**). Each group was then challenged with *P. berghei* and oocyst numbers were examined on Day 8 post-infection. (**D**) Before infection, the effects of clodronate treatment on the percentage of granulocytes was examined in the presence or absence of rTNF-α to confirm our experimental approach. (**E**) Infection outcomes following granulocyte depletion and rTNF-α treatment, with oocyst numbers and infection prevalence evaluated at 8 days post infection. For both **D** and **E**, “+” denotes treatment with rTNF-α, while “-” indicates treatment with 1X-PBS. For all experiments, the dots represent the respective measurements from an individual mosquito. The red horizontal lines represent the median oocysts numbers, while infection prevalence (% infected/total) is depicted as chart pies below each figure containing infection data. Data were combined from three or more independent experiments. Statistical significance was determined using either Mann-Whitney (individual comparisons) or Kruskal-Wallis with a Dunn’s multiple comparison test (multiple comparisons) to assess oocyst numbers, the number of attached hemocytes, or the percentage of granulocytes. A Fisher’s Exact test was performed to measure differences in infection prevalence. Asterisks denote statistical significance (* *P* < 0.05, ** *P* < 0.01, *** *P* < 0.001, **** *P* < 0.0001). ns, not significant; n= numbers of individual mosquitoes examined.

**Figure 6.**
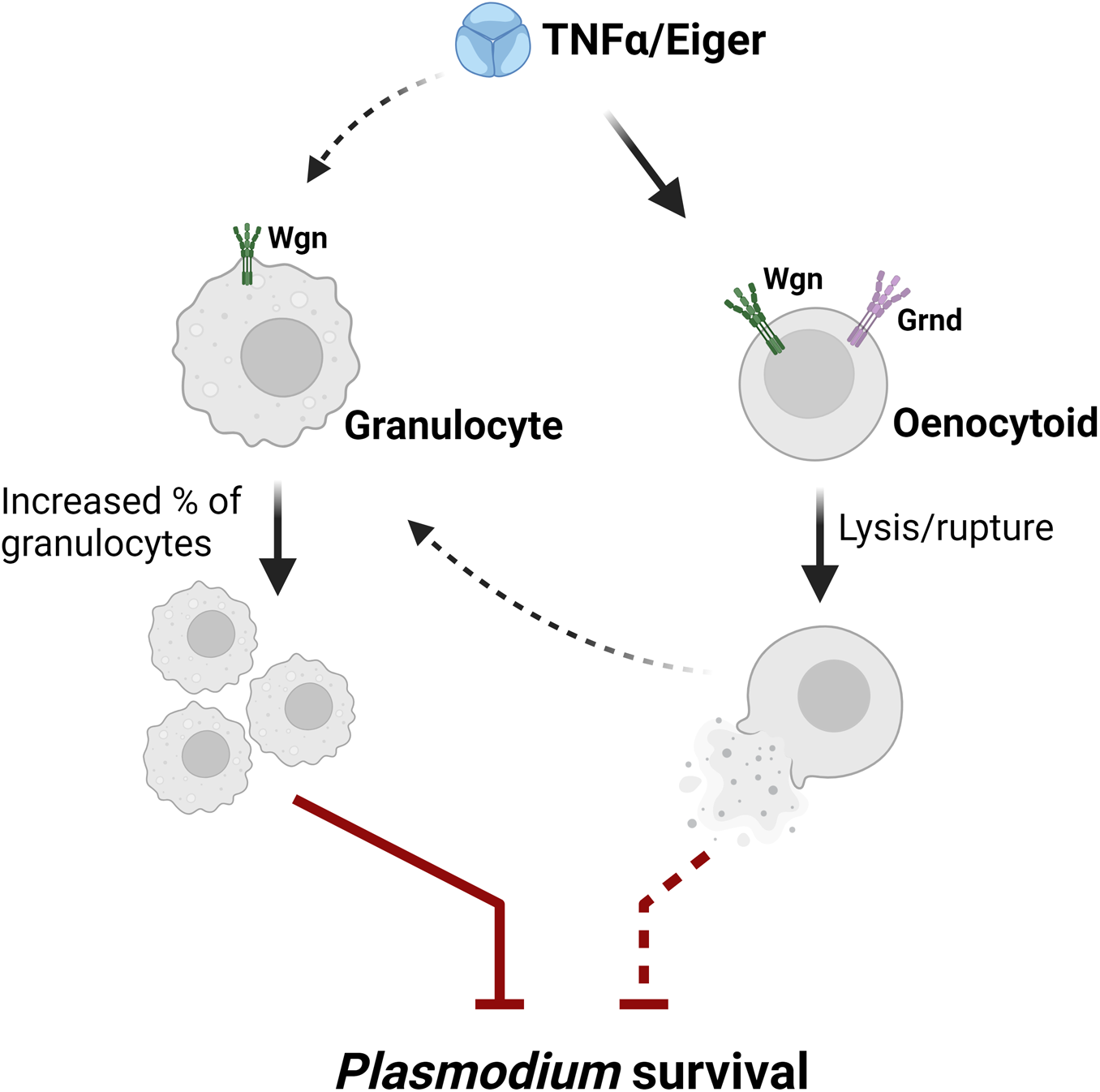
Proposed model of TNF signaling on mosquito immune cells and their contributions to anti-*Plasmodium* immunity.

## Discussion

Mosquito innate immunity is an integral component of vector competence [53], therefore understanding the immune mechanisms that influence *Plasmodium* survival is essential for ongoing efforts to limit malaria transmission. While several conserved immune signaling pathways, such as Toll, IMD, and JAK/STAT, have been previously implicated in mosquito vector competence [54–57], here we provide direct evidence for the role of TNF signaling in mosquito immune function.

Our expression analysis of the mosquito TNF signaling components, Eiger, Wgn, and Grnd, suggests that TNF signaling is ubiquitous across mosquito tissues, with expression detected across midgut, hemocyte, and fat body tissues. While the expression of the receptors remained relatively constant across physiological conditions, the mosquito TNF ortholog *Eiger* was more responsive to blood-feeding or *P. berghei* infection, displaying significant induction in hemocytes. While this implies that TNF signaling is further amplified by mosquito hemocytes, potential post-translational modifications, which require cleavage to release TNF-α/Eiger in its active soluble form [58], complicate our ability to make broad sweeping conclusions from these data. However, TNF signaling has been previously implicated in *Drosophila* as a master regulator of midgut homeostasis regulating lipid metabolism and intestinal stem cell (ISC) proliferation to maintain tissue integrity [48]. In addition, *Eiger* is expressed in *Drosophila* hemocytes in response to the midgut-derived reactive oxygen species (ROS), which ultimately triggers ISC proliferation [39]. Since mosquito blood-feeding represents a significant physiological event causing distention and damage to the midgut epithelium, the repair of the midgut epithelium may require similar roles of intestinal stem cell differentiation [59]. This is supported by previous studies that highlight the involvement of stem cells in mosquito midgut homeostasis in response to blood-feeding, oxidative stress, and infection [60,61]. Considering the function of Eiger/Wgn signaling in *Drosophila* epithelial turnover [39,40], there is potential that the observed increase in Eiger expression following blood-feeding and infection could similarly stimulate ISC proliferation to promote midgut homeostasis, a process that may potentially involve hemocyte function. However, at present, direct roles of TNF signaling in mosquito midgut regeneration have yet to be determined.

Using the paired approach of rTNF-α injection and RNA interference (RNAi) to address the potential roles of mosquito TNF signaling, we demonstrate that TNF signaling is an integral component of anti-*Plasmodium* immunity in *Anopheles gambiae*. While rTNF-α injection (and presumably overexpression of the pathway) results in reduced malaria parasite survival, loss of *Eiger*, *Wgn*, or *Grnd* prior to *P. berghei* challenge each cause an increase in *Plasmodium* oocyst numbers. While in vertebrates, TNF signaling can initiate distinct cellular responses mediated by TNFR1 and TNFR2 [62], our double-knockdown experiments suggest that both *Wgn* and *Grnd* are required to initiate anti-*Plasmodium* immunity. At present, it is unclear if Wgn and Grnd act as a heterodimeric receptor to promote TNF signaling, despite the lack of support from other systems. Alternatively, based on recent studies [63], Wgn and Grnd may differ in their subcellular localization and functional roles in the processing of TNF-mediated signals.

Although TNF-α is a well-established proinflammatory cytokine known for its role in regulating various aspects of macrophage function in vertebrates [64,65], our understanding of TNF signaling on insect immune cells has been limited. Previous studies have implicated Eiger/TNF in phagocytosis and host survival to pathogen infection [44–46], supporting the conservation of the TNF signaling pathway across insect taxa. Moreover, TNF signaling has been implicated in regulating phagocytic immune cell populations in solitary locusts [46] and in promoting crystal cell rupture in *Drosophila* [42]. Here, we provide evidence that TNF signaling similarly regulates mosquito immune cell function by increasing the percentage of circulating granulocyte populations and in driving oenocytoid immune cell lysis.

As important immune sentinels, granulocytes are the primary phagocytic immune cells in the mosquito, either circulating in the open hemolymph or attached to various mosquito tissues [20]. While data support the increase in the percentage of granulocytes following rTNF-α treatment, our limited understanding of mosquito hemocyte biology and lack of genetic tools makes this a challenging phenotype to address. As a result, it remains unclear if TNF signaling via Wgn/Grnd promotes differences in cell adherence (from sessile cells to in circulation), granulocyte activation, or the differentiation of precursor cells to granulocytes. This is further complicated by the requirement of *Grnd* for the rTNF-α-mediated effects on granulocyte populations, which based on the data presented here and previous single-cell studies [22], suggest that *Grnd* is not expressed in granulocytes. As a result, we speculate that the TNF-mediated increase in granulocytes is indirect, and potentially caused by the release of other molecules resulting from oenocytoid lysis or the production from other tissues such as the fat body.

Similar to previous studies in *Drosophila* demonstrating the role of *Eiger* in crystal cell lysis [42], we demonstrate that rTNF-α promotes the lysis/rupture of mosquito oenocytoids, the equivalent of *Drosophila* crystal cells. With evidence that *Eiger* is required for the release of prophenoloxidase [42] and melanization activity [43], our data displaying increased PO-activity following rTNF-α injection provide support for a similar mechanism in mosquitoes via TNF signaling. These results are further supported by complementary studies where we have previously shown that oenocytoid lysis is triggered by prostaglandin E2 (PGE2) in a concentration-dependent manner [19]. Similar to our previous observations [19], TNF-α reduced the expression of PPO1, PPO8, and PGE2R, genes enriched in oenocytoid populations [22], while having no effect on the expression of the phagocytic granulocyte marker *eater* [17,22]. Taken together, our data suggest that TNF signaling regulates oenocytoid rupture, leading to the release of their cellular contents in the hemolymph. This includes the release of prophenoloxidases, which have been previously implicated in mosquito anti-bacterial [19] and anti-*Plasmodium* immunity [17,19].

Previous studies have implicated both granulocytes and oenocytoids in limiting malaria parasite survival in the mosquito host [16,17,19,66]. However, with both immune cell subtypes displaying phenotypes associated with TNF signaling, we sought to further examine the potential roles of mosquito hemocytes in TNF-mediated parasite killing. Evidence suggests that *Plasmodium* killing in mosquitoes is multimodal with distinct immune responses that target either invading ookinetes or immature oocysts [52,53]. When we examined early (day 2) oocyst numbers as a proxy to determine the success of ookinete invasion [15,67], early oocyst numbers were significantly reduced in rTNF-α-injected mosquitoes, suggesting that TNF signaling contributes to *Plasmodium* ookinete killing. Given the importance of granulocytes in mediating ookinete recognition by mosquito complement [16,17], this suggests that the TNF regulation of granulocyte function is central to early-phase immune responses targeting the ookinete. This is further supported by our observations of increased hemocyte attachment to the midgut following rTNF-α treatment. Through the use of clodronate liposomes to deplete phagocytic granulocyte populations [17,22,50,51], we confirm the involvement of granulocytes in TNF-mediated parasite killing. Yet, the incomplete effects of granulocyte depletion to abrogate parasite killing following rTNF-α treatment suggests that additional components contribute to limiting parasite survival. However, it is unclear if these TNF-mediated responses are produced by granulocyte populations not influenced by clodronate depletion [17] or if these immune responses are produced by other immune cell subtypes or tissues. With observations that rTNF-α also influences oenocytoid rupture and PO activity, which are similar to the late-phase immune responses limiting oocyst survival via prostaglandin signaling [19], we propose a model based on which the influence of TNF signaling on anti-*Plasmodium* immunity is mediated by both granulocyte and oenocytoid immune functions (summarized in **Fig 6**). Alternatively, we cannot exclude the potential that TNF-mediated parasite killing is mediated in part by humoral responses produced by the fat body, yet due to the systemic nature of RNAi, we currently lack the genetic tools to examine the cell-or tissue-specific contributions of TNF signaling in *An. gambiae*.

In summary, our findings provide important new insights into the roles of TNF signaling in *An. gambiae,* demonstrating the effects of TNF signaling in limiting malaria parasite survival and immune cell regulation. While further study is required to fully determine the influence of TNF signaling on mosquito physiology and immune function, there is significant evidence, in addition to that provided herein, that TNF signaling is a central component that defines mosquito vectorial capacity and the susceptibility to *Plasmodium* infection in natural mosquito populations [36]. As a result, our study represents an important contribution to our understanding of the mechanisms of malaria parasite killing and the collective efforts to develop novel approaches for malaria control.

## Materials and Methods

### Ethics Statement

All protocols and experimental procedures regarding vertebrate animal use were approved by the Animal Care and Use Committee at Iowa State University (IACUC-18-228).

### Mosquito Rearing and *Plasmodium* Infection

*Anopheles gambiae* mosquitoes (Keele strain) were reared at 27°C and 80% relative humidity, with a 14:10 h light: dark photoperiod cycle. Larvae were fed on commercialized fish flakes (Tetramin, Tetra), while adults were maintained on a 10% sucrose solution and fed on commercial sheep blood (Hemostat) for egg production.

Female Swiss Webster mice were used for mosquito blood-feeding and infections with a *Plasmodium berghei* (*P. berghei*) transgenic strain expressing mCherry [15,17]. Following infection, mosquitoes were incubated at 19°C for either two days or eight days before individual mosquito midguts were dissected to determine parasite loads by examining oocyst numbers under a compound fluorescent microscope (Nikon Eclipse 50i; Nikon).

### RNA extraction and gene expression analyses

Total RNA was extracted from whole mosquito samples or dissected tissues using Trizol (Invitrogen, Carlsbad, CA). RNA from perfused hemolymph samples was isolated using the Direct-Zol RNA miniprep kit (Zymo Research). Two micrograms of non-hemolymph-derived or 200ng of hemolymph-derived total RNA were used for first-strand synthesis with the RevertAid reverse transcriptase kit (Thermo Fisher Scientific). Gene expression analysis was performed with quantitative real-time PCR (qPCR) using PowerUp SYBRGreen Master Mix (Thermo Fisher Scientific) as previously described [17]. qPCR results were calculated using the 2^−ΔCt^ formula and standardized by subtracting the Ct values of the target genes from the Ct values of the internal reference, *rpS7*. All primers used in this study are summarized in **Table S1**.

### Gene identification and silencing

The mosquito orthologs of known *Drosophila* TNF-α signaling components, Eiger (AGAP006771), Wengen (AGAP000728), and Grindelwald (AGAP008399), were identified using the OrthoDB database [68]. To address gene function, T7 primers specific to each candidate gene (**Table S1**) were used to amplify DNA templates from whole female mosquito cDNA for dsRNA production and RNAi as previously [15,17]. PCR products were purified using the DNA Clean & Concentrator kit (Zymo Research) following gel electrophoresis to test for target specificity. dsRNA synthesis was performed using the MEGAscript RNAi kit (Thermo Fisher Scientific), with the concentration of the resulting dsRNA adjusted to 3μg/μl. For RNAi, adult female mosquitoes (3-5 days old) were anesthetized on a cold block and injected intrathoracically with 69nl of dsRNA targeting each gene or GFP as a negative control. Co-silencing of *Wgn* and *Grnd* was accomplished by injecting mosquitoes with a solution consisting of equal parts of the dsRNA suspensions targeting each gene. Injections were performed using Nanoject III manual injector (Drummond Scientific). To assess gene-silencing, groups of ten mosquitoes were used to analyze the efficiency of dsRNA-mediated silencing at 2 days post-injection via qPCR.

### Injection of human recombinant TNF-α

Recombinant human TNF-a (rTNF-α; Sigma #H8916) was resuspended in 1X PBS to a stock solution of 0.72ng/nl. Naive or dsRNA-injected mosquitoes were anesthetized and intrathoracically injected with either 69nl of 1X PBS (control) or the stock solution of rTNF-α to administer 50ng of protein per individual. Following injection, mosquitoes were maintained at 27^°^C for 24hrs then used for downstream infection experiments or hemocyte analysis.

### Hemocyte counting

Mosquito hemolymph was collected by perfusion using an anticoagulant buffer of 60% v/v Schneider’s Insect medium, 10% v/v Fetal Bovine Serum, and 30% v/v citrate buffer (98 mM NaOH, 186 mM NaCl, 1.7 mM EDTA, and 41 mM citric acid; buffer pH 4.5) as previously described [15,17,66]. For perfusions, incisions were performed on the posterior abdomen, then anticoagulant buffer (∼10μl) was injected into the thorax. Collected perfusate from individual mosquitoes were placed in a Neubauer Improved hemocytometer and observed under a light microscope (Nikon Eclipse 50i; Nikon) to distinguish hemocyte subtypes by morphology and determine the proportion of granulocytes out of the total number hemocytes in the sample.

### Hemocyte gene expression analysis

Following mosquito injections with rTNF-α, perfused hemolymph from at least 20 mosquitoes was used for RNA extraction, cDNA synthesis, and qPCR to estimate the expression levels of hemocyte subtype gene markers (PPO1, PPO8, and PGE2R; oenocytoid-specific, and Eater; granulocyte-specific) [19,22].

### Characterization of hemolymph PO activity

To determine the effects of TNF-α on phenoloxidase (PO) activity, naïve mosquitoes were injected with either 1X PBS (control) or rTNF-α. At 24h post-injection, hemolymph was perfused from 15 mosquitoes using nuclease-free water as previously described [17,19,69]. The perfusate (10μl) was added to a 90μl suspension of 3, 4-Dihydroxy-L-phenylalanine (L-DOPA, 4 mg/ml), then incubated at room temperature for 10 min prior to measurements of PO activity using a microplate reader at 490nm. Samples were measured using six independent measurements at 5 min intervals.

### *Plasmodium* infections following rTNF-α injection

After the injection of rTNF-α as described above, both control and experimental groups were challenged on a *P. berghei*-infected mouse. After selecting for blood-fed mosquitoes on ice, mosquitoes were kept at 19^°^C until oocyst survival was assessed at either 2- or 8-days post-infection.

To determine the TNF-α-mediated responses against *Plasmodium* in a granulocyte-depleted background, 3-day-old mosquitoes were first injected with either control or clodronate liposomes as previously [17]. At 24h post-injection, each group was treated with 50ng of rTNF-α or 1X PBS as control. Following an additional 24h incubation, surviving mosquitoes were challenged with *P. berghei*, with oocyst numbers examined from dissected midguts at either 2- or 8-days post-infection.

### Analysis of *Wgn* and *Grnd* expression in hemocyte subtypes

To determine the expression of TNF signaling components in mosquito hemocyte subpopulations, the expression of *Wgn* and *Grnd* was referenced with previous single-cell transcriptomic data for *An. gambiae* hemocytes [22]. Further validation was performed using methods of granulocyte depletion via clodronate liposomes as described above [17] to confirm the presence/absence of *Wgn* and *Grnd* expression in granulocyte populations using qPCR.

### Immunofluorescent analysis of hemocytes attached to midguts

Hemocyte attachment to mosquito midguts in response to rTNF-α treatment was examined by immunofluorescence analysis as previously described [18] with slight modification. Two days after treatment with either rTNF-α or 1XPBS, mosquitoes were injected with 69nl of 100μM Vybrant CM-DiI cell labeling solution (ThermoFisher) and allowed to recover for 30min at 27°C. Mosquitoes were injected with 200nl of 16% paraformaldehyde (PFA), then the entire mosquito was immediately submerged/ incubated in a solution of 4% PFA for 40 sec prior to transfer in ice-cold 1XPBS for midgut dissection. Dissected midguts were incubated overnight in 4% PFA at 4°C for fixation. The following day, midguts were washed with ice-cold 1XPBS three times and permeabilized with 0.1% TritonX-100 for 10 minutes at room temperature. After washing three times with 1XPBS, tissues were blocked with 1% BSA in 1XPBS for 40 minutes at room temperature and stained with Phalloidin-iFluor 405 Reagent (1:400 in PBS; abcam, ab176752) for 1 hour to visualize actin filaments. Midguts were washed with 1XPBS to remove excess staining, placed on microscope slides and mounted with ProLong Diamond Antifade Mountant (ThermoFisher). Samples were imaged by fluorescence microscopy using a Zeiss Axio Imager 2 and analyzed to determine the number of hemocytes attached to individual midguts from each experimental condition.

## Supporting information

Supporting Information containing Figures S1 to S2 and Table S1

## Acknowledgements

This work was supported by R21AI144705 and R21AI166857 to RCS from the National Institutes of Health, National Institute of Allergy and Infectious Diseases.

