## Supporting Information containing Figures S1 to S2 and Table S1 for "TNF signaling mediates cellular immune function and promotes malaria parasite killing in the mosquito *Anopheles gambiae*"

### Included supporting information

#### *Supplemental Figures*

**Figure S1.** Silencing efficiency of *Eiger*, *Grnd*, and *Wgn* genes in *An. gambiae.*

**Figure S2.** Expression of *Eiger*, *Wgn*, and *Grnd* in mosquito hemocyte populations.

#### *Supplemental Tables*

**Table S1.** List of primers used for gene expression and RNAi.

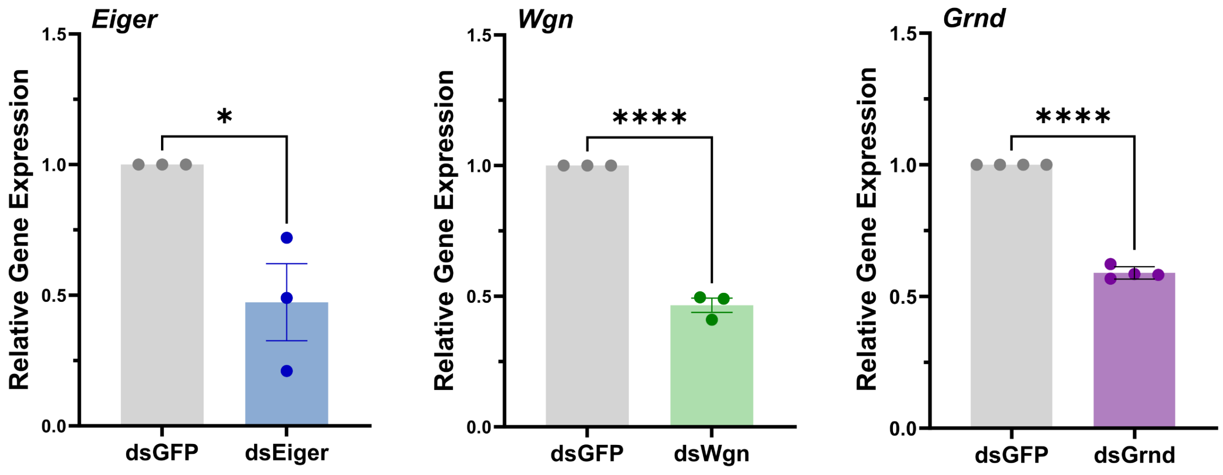

**Figure S1. Silencing efficiency of *Eiger*, *Grnd*, and *Wgn* genes in *An. gambiae.*** Naïve adult female mosquitoes were injected with dsRNA targeting *GFP* (control), *Eiger*, *Wgn*, or *Grnd*. Two days post-injection, whole-body mosquitoes (10-15 total) were collected for RNA extraction, followed by gene expression analysis using qPCR. Data from three or more independent experiment were examined for statistical significance using an unpaired student's t-test. Asterisks indicate significance (* $P < 0.05$, **** $P < 0.0001$).

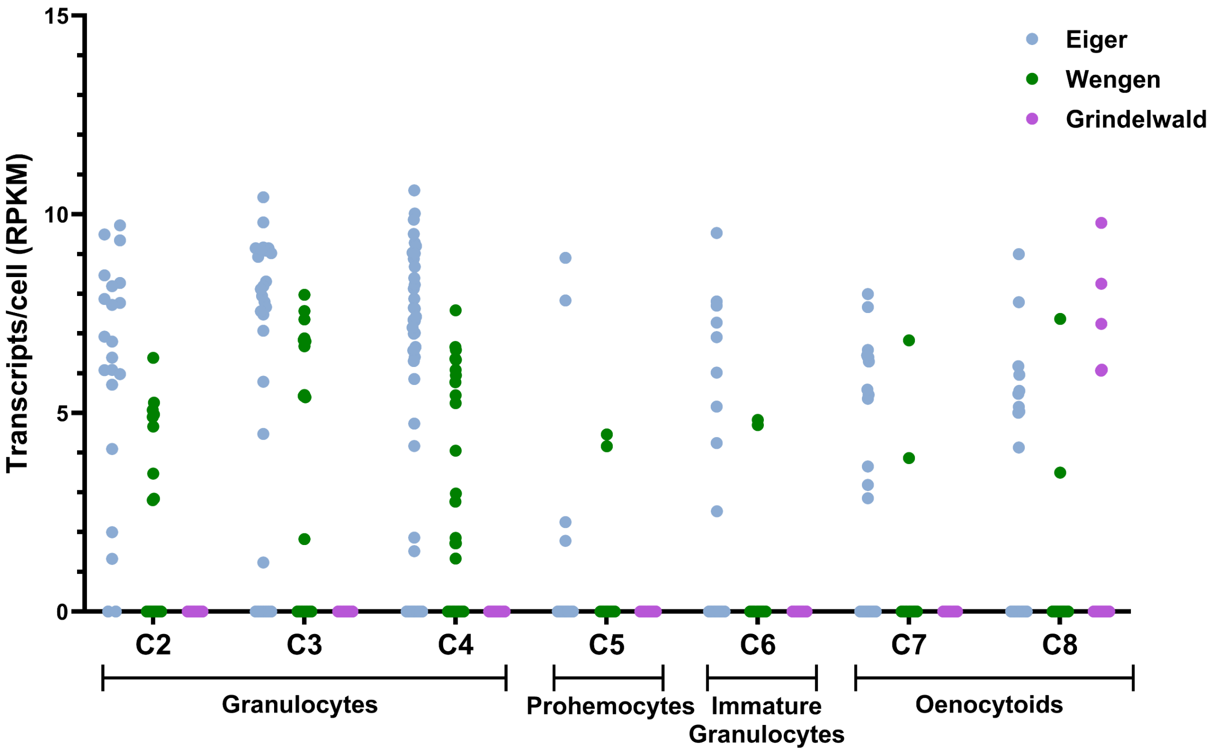

**Figure S2. Expression of *Eiger*, *Wgn,* and *Grnd* in mosquito hemocyte populations.** Using previously published single-cell RNA-seq data of *An. gambiae* hemocytes [22], the expression of Eiger, Wgn, and Grnd was evaluated for each mosquito immune cell subtype. Each dot represents a cell-specific gene expression value. RPKM, Reads Per Kilobase Per Million.

**Table S1. List of primers used for gene expression and RNAi.** Small letters indicate the T7 promoter sequence.

| Gene | Forward (5'-3') | Reverse (5'-3') |
|---|---|---|
| AgGrnd-qPCR | CAAGGCGGTGCCGAAGAATG | GCTTTCCGTCGAATCTTCCG |
| AgGrnd-T7 | taatacgactcactatagggGTGCGTGTGTGTGTGTTCAG | taatacgactcactatagggCGGACCCTGTTCCTTCTTGT |
| AgEiger-qPCR | TCCGCTGGGATGTAGAAAATCG | GGCGTGGTGGTGCTGTGATG |
| AgEiger-T7 | taatacgactcactatagggCAATGAGCTGAACGCTGGAA | taatacgactcactatagggGGCTCGTTGATGGTAAGCTG |
| AgWgn-qPCR | GAGGAGATCCTGTGGGACTG | GCGTCAAAGTGCTTCTCGAT |
| AgWgn-T7 | taatacgactcactatagggAAGCATTATCGGCCCAGCTC | taatacgactcactatagggGGGAAGCCAGATTTTGGATCTAG |
